# Temporal stability and niche partitioning of bacterial communities in paired residential sink P-traps

**DOI:** 10.64898/2026.02.17.706431

**Authors:** Megan S. Hill, Will Stiffler, Jorden T. Rabasco, Ivory C. Blakley, Shan Sun, Anthony A. Fodor, Claudia K. Gunsch

## Abstract

Sink P-traps harbor microbial communities derived from environmental and human sources, yet longitudinal studies examining their stability and assembly dynamics remain limited. Here, we present, to our knowledge, the longest continuous characterization of bacteria collected from residential sink P-traps, with daily sampling over two months (n = 61 days). Samples were collected from paired sinks in a shared bathroom to identify dominant taxa, quantify temporal stability, and determine how occupant usage patterns influence community assembly. Using full-length 16S rRNA gene sequencing, we identified 3,865 unique taxa, with both sinks dominated by common sink-associated genera, including *Pseudomonas*, *Citrobacter*, *Klebsiella*, and *Arcobacter*. Despite sharing identical plumbing, environmental conditions, and cleaning regimes, the two sinks maintained distinct communities (p < 0.001). Temporal stability analyses revealed notable differences: Sink A (male; hand washing, toothbrushing, shaving) exhibited deterministic dynamics with low variability (CV = 4.9%), significant temporal autocorrelation (p = 0.001), and predictable trajectories, with time explaining 49.9% of community variation. In contrast, Sink B (female; hand washing, toothbrushing, face washing, mouthwash use) displayed stochastic dynamics with high volatility (CV = 26.5%), no significant autocorrelation (p = 0.53), and minimal temporal predictability. Differential abundance analysis revealed that Sink B was enriched in anaerobes, biofilm-forming taxa, oral microbiome associates, and preservative-resistant and lipid-degrading bacteria, while Sink A harbored a more aerobic, skin-associated community. These findings demonstrate that individual usage patterns (particularly exposure to biocidal agents) can alter P-trap community structure and temporal dynamics, with implications for microbial community prediction in residential and healthcare settings.

**Importance:** Sink drains are increasingly recognized as reservoirs for antimicrobial-resistant pathogens, yet we lack fundamental knowledge about what drives bacterial community dynamics in these environments. By sampling paired residential sink P-traps daily for two months, we show that individual-specific behaviors, such as using products with a biocidal effect, can alter community composition from a stable, predictable state to one characterized by stochastic fluctuations. The sink exposed to mouthwash and face wash harbored more anaerobes, biofilm formers, and oral bacteria, suggesting that repeated exposure promotes disturbance-tolerant taxa rather than reducing bacterial colonization. These results provide a baseline ecological framework for understanding P-trap microbiomes and suggest that predictive monitoring of sink-associated pathogens might be feasible in stable environments but more difficult when there is variable antimicrobial exposure -- a finding directly relevant to hospital infection control.

## Introduction

Sink P-traps host microbial communities derived from both environmental and human-associated sources, including water, personal care products, and skin and oral microbiota (1–3). The selective conditions of this microenvironment promote colonization and growth of bacteria adapted to oligotrophic, chemically variable, and intermittently aerobic conditions, including species within the *Pseudomonas*, *Sphingomonas*, and *Mycobacterium* genera (4, 5). P-traps are continually wet with periodic hot water flushing, nutrient pulses, and exposure to surfactants and disinfectants, selecting for stress tolerant bacteria and biofilm-formers (6). These biofilms can promote the persistence of potential pathogens and act as a reservoir that enables microbial reseeding of upstream plumbing components, room surfaces, and indoor bioaerosols, increasing the risk of pathogen exposure in healthcare settings (7–10). This is of high concern, as hospital-acquired infections (HAIs) are contracted by 1 in every 31 patients in the US (11, 12). Therefore, it is imperative that we understand sink microbial ecology, the relative impact of differential usage between individuals, and the stability and predictability of bacterial communities over short and long temporal scales.

P-trap studies have largely focused on species of concern (e.g., pathogens or material-degrading microbes) or relied on sparse or cross-sectional study designs, and microbial community stability and assembly dynamics in hospital sinks is inherently difficult to assess. Individual hospital sinks might be used by multiple people per day, with usage patterns fluctuating based on patient occupancy and diagnosis, staff behavior, and clinical workflows. These sinks are cleaned frequently with disinfectants and cleaning agents, and there is continual deposition from individuals colonized by or infected with pathogens. As a result, hospital sink microbiomes experience intense and irregular disturbance events, making it challenging to disentangle the relative contributions of stochastic dispersal, environmental filtering, and biological interactions on community structure. In contrast, residential homes can provide a tractable framework for understanding the relative roles of disturbance, selection, and dispersal in sink-associated microbiomes. Insights gained from residential P-traps can inform hospital sink studies by establishing baseline expectations for community stability, resilience, and succession in the absence of extreme disturbance and high human traffic. More broadly, residential sink microbiomes might inform ecological principles that govern pathogen persistence and community resistance, ultimately guiding the development of more effective sink design, cleaning strategies, and microbial management interventions in healthcare environments.

Here, we present, to our knowledge, the longest continuous characterization of bacterial communities in residential sink P-traps to date, to resolve short-term ecological dynamics under real-world household use. We collected daily samples over a two-month period (n = 61 consecutive days) from paired sink P-traps within a shared residential bathroom that were used by different occupants (1 male and 1 female). Additionally, none of the shared personal care or cleaning items contained antimicrobials, while one person used nightly mouthwash that acts as a biocidal agent. This design allows us to examine community structure, temporal stability, and inter-sink variability, based on differential usage patterns and individual exposures, while controlling for a range of factors that could potentially alter bacterial colonization and growth (e.g., home and sink characteristics, ambient temperature and humidity, cleaning frequency, and common personal care products). By resolving day-to-day microbial dynamics in residential sinks, this study aims to (i) identify dominant bacterial taxa in household sink P-traps, (ii) quantify the stability and variability of P-trap microbiomes at a high temporal resolution over a long period of time, (iii) identify factors that alter community assembly dynamics based on occupant usage patterns, and (iv) provide an ecological framework for understanding bacterial persistence and niche partitioning that can be applied to other built environment types.

## Materials and Methods

Bacteria were collected from two bathroom sink P-traps within a residential home in Chapel Hill, North Carolina, daily for two months (n = 61 days) during May–June 2024. The sinks were located in the same bathroom and were used by different occupants throughout the duration of the study. Both sinks were subject to the same cleaning regimes (e.g., frequency, cleaning product), and both occupants used the same brand of toothpaste and hand soap – none of these products contained antimicrobial compounds. Both sinks were used daily for routine hand washing and toothbrushing. However, Sink A (male) was additionally used for daily shaving, and Sink B (female) was used for nightly face washing and mouthwash use.

### Sample Collection, Processing, and Sequencing

Samples were collected using a sterile tubing and syringe assembly connected via a T-adapter. The tubing was inserted into the P-trap, and biofilm and liquid were agitated by repeatedly filling and expelling the syringe ten times. Approximately 43 mL of each sample was collected into sterile 50 mL centrifuge tubes and stored at 4 °C until processing, and sample tubes were labeled with the date, sink identifier, and a three-digit code denoting the approximate abundance of visible biofilm fragments (> 5 mm, > 1 mm, and < 1 mm in diameter). Negative control samples were collected every seventh day by pipetting sterile deionized water from the bathroom countertop adjacent to the sink, and an additional P-trap sample was collected at each control time point and preserved in 20% glycerol for archival storage.

P-trap samples were filtered through 0.22 µm polyethersulfone (PES) membranes using a sterile Büchner funnel, with the filtrate passed at ∼1 mL/sec and allowed to drain for an additional minute after completion, with a sterile control included in each filtration batch. Filters designated for downstream molecular analysis were transferred into 1.5 mL microcentrifuge tubes, and DNA was extracted using the DNeasy PowerSoil Pro Kit (Qiagen, Valencia, CA, USA) following manufacturer instructions, with minor modifications. Briefly, filters were transferred to sterile petri dishes, cut into 5 mm pieces using a flame-sterilized scalpel and tweezers, and placed into PowerBead tubes for mechanical disruption using a bead beater (30 sec on, 30 sec off, 30 sec on), Extracted DNA was stored at −20 °C prior to downstream processing and sequencing of the full-length 16S rRNA gene.

### Sequencing and Data Analysis

Extracted DNA was quantified using a Qubit fluorometer and normalized to 1 ng/µl. Two nanograms of total DNA per sample were amplified using the PacBio Kinnex 16S kit (PacBio, Menlo Park, California, USA) with Phusion Plus PCR Master Mix (Thermo Fisher Scientific, Waltham, MA, USA) and the universal 27F/1492R primer pair (13); each at a final concentration of 0.3 µM and containing unique barcodes and Kinnex adaptors. PCR conditions were: initial denaturation at 98 °C for 30 s; 25 cycles of 98 °C for 10 s, 57 °C for 20 s, and 72 °C for 75 s; and a final extension at 72 °C for 5 min.

Amplification success and amplicon size (∼1,500 bp) were verified on an E-Gel (Invitrogen). Amplicons were pooled without cleanup in volumes (1–20 µl per reaction), based on gel band intensity, and purified and concentrated using 1.1× SMRTbell® cleanup beads (PacBio, Menlo Park, California, USA). The pooled library was eluted in 50 µl of Low TE buffer, quantified via Qubit, and stored at −20 °C prior to Kinnex PCR for concatenation and circularization. Kinnex PCR, size selection, and final cleanup were performed according to the manufacturer’s protocol (PacBio). Libraries were loaded onto a PacBio SMRT® Cell and sequenced on the Revio system at the Duke Sequencing and Genomic Technologies Shared Resource.

Raw sequencing data was processed into amplicons via standard DADA2 processes (version 1.37.0) (14). Briefly, the primers were removed and the lengths of the reads were trimmed to 1600 bp. These raw reads were then resolved into amplicons using a binned quality score error model. The pseudopooling method was then employed to resolve the reads into the amplicon sequence variants (ASVs). The ASVs were then taxonomically assigned via the assignTaxonomy() function within DADA2, using the SILVA database (silva_nr99_v138.1_wSpecies_train_set) (15). With the exception of the differential abundance analysis (for which the unrarefied data were used), samples were then rarefied to 14,700 reads per sample (loss of n = 9 samples) and transformed using a centered log-ratio (CLR) transformation. Data were analyzed in an R environment (v. 4.3) and visualized with ggplot2 (16).

Differences in Chao1 and Shannon diversity indices were compared using the phyloseq package (17). Overall differences between sinks were calculated with Wilcoxon signed-rank tests (18), and alpha diversity among weeks was tested using Kruskal-Wallis tests, with Dunn’s post-hoc test (Bonferroni-corrected) used to identify pairwise differences (19). Taxa relative abundances were calculated using mctoolsr (20). Differences in community composition were calculated with Aitchison distance, and compared with a permutational multivariate analysis of variance (PERMANOVA) test (21) using the vegan package (22).

To assess the temporal stability and predictability of sink bacterial communities, differences in beta diversity (Bray-Curtis dissimilarity) were calculated between the initial sampling timepoint and all other samples for each sink. Multiple complementary approaches were then used to characterize community dynamics over the sampling period. The coefficient of variation (CV) of beta diversity values were calculated to quantify relative stability, where lower CV indicates more consistent community composition over time. The mean squared successive difference (MSSD) was computed to measure trajectory volatility, with lower values indicating smoother temporal dynamics (23). Temporal autocorrelation was assessed using the autocorrelation function (ACF) at lags 1–15, to determine whether community states at one timepoint predicted states at subsequent timepoints. The Ljung-Box test was used to formally test for significant autocorrelation structure in the time series (24). Generalized additive models (GAMs) were fit to model non-linear temporal trends with a smooth term for time using restricted maximum likelihood (REML) estimation (25). The proportion of deviance explained was used to assess how well temporal dynamics could be captured by the model, with significance of the smooth term indicating whether time was a meaningful predictor of community state. Autoregressive integrated moving average (ARIMA) models were fit using automatic model selection based on AIC (26). Model performance was evaluated using root mean squared error (RMSE) and mean absolute error (MAE) on the training data.

To then assess true predictive performance, a rolling-window cross-validation scheme was used where models were trained on the first 70% of timepoints and iteratively used to predict each subsequent timepoint. Cross-validation RMSE was calculated as the primary measure of out-of-sample predictive accuracy. To determine the minimum sampling frequency required to accurately characterize temporal dynamics, the daily time series was systematically subsampled at intervals of 2, 3, 5, 7, and 14 days. For each interval, 10 random starting offsets were applied and results averaged to minimize subsampling bias. At each frequency, the CV, MSSD, GAM deviance explained, lag-1 autocorrelation with the Ljung-Box test, and leave-future-out cross-validation RMSE were recalculated and compared to the daily reference values. All temporal analyses were performed using the mgcv package v1.9 for GAMs (25) and the forecast package v8.21 for autocorrelation and ARIMA modeling (26).

Finally, to better understand taxonomic-level differences, a differential abundance analysis was conducted, using ALDEx2 (27). To reduce false positives, taxa with an abundance of less than 1e-5% were removed prior to analysis. The model was run with the ‘decom = iqlr’ flag, and taxa with a p-value < 0.05 were retained. After characterization, the top 15 differentially abundant taxa (i.e., those with the greatest effect size) were then annotated, based on phenotype and/or source.

## Results

Overall, we identified 3865 unique taxa, with similar richness (∼735–745 ASVs) and evenness (∼4.0 Shannon) between the two sinks (Chao1: p = 0.864, Shannon: p = 0.760). Daily fluctuations in alpha diversity were observed (Fig. S1), with significant week-to-week variation (Fig. 1a; Kruskal-Wallis: Sink A, Chao1 p < 0.001, Shannon p = 0.027; Sink B, Chao1 p = 0.037, Shannon p = 0.033). However, both sinks were dominated by similar taxa (Fig. 1b), including common sink-associated groups, such as *Pseudomonas* spp. (5), *Citrobacter freundii* (28), *Klebsiella* spp. (29, 30), and *Arcobacter butzleri* (31). Despite these similarities in abundant bacterial groups, each sink maintained a distinct bacterial community (Fig. 1c; Aitchison Distance: p < 0.001) – a differentiation that was also observed at higher levels of taxonomic classification (Fig. S2; p ≤ 0.001 for all comparisons; effect sizes: species R^2^ = 0.278, genus R^2^ = 0.208, family R^2^ = 0.135, order R^2^ = 0.110).

**Figure 1.**
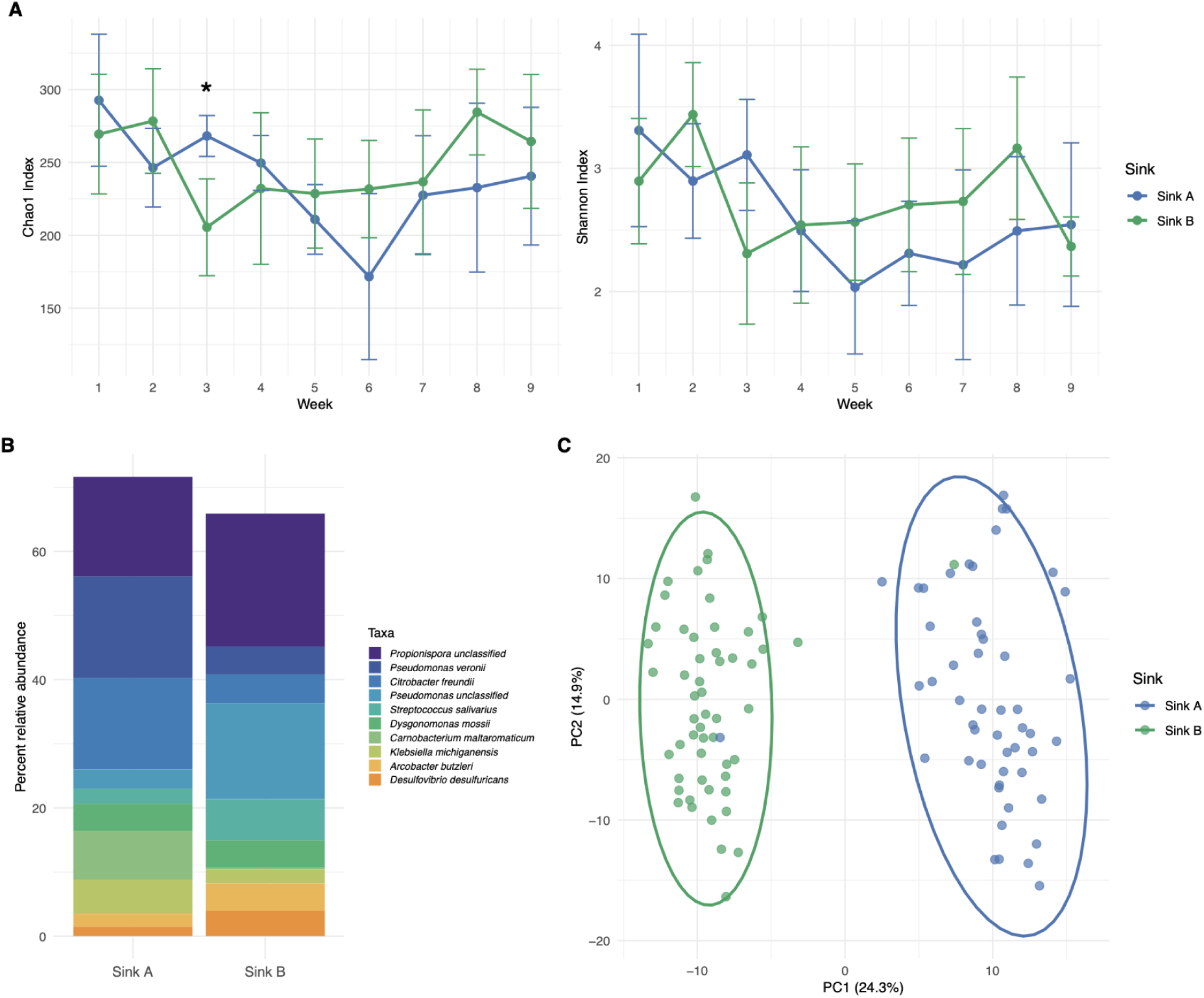
Sink microbial diversity and community structure over time. (A) Alpha diversity metrics (Chao1 richness and Shannon diversity index) for Sink A (blue) and Sink B (green) across nine weeks. Points represent mean values with standard deviation error bars. Asterisks denote significant differences between sinks (Wilcoxon rank-sum test, p < 0.05). (B) Relative abundance of the top 10 most abundant bacterial taxa across all samples, based on sink. (C) Principal coordinates analysis (PCoA), as calculated based on Aitchison distance. Ellipses represent 95% confidence intervals.

Temporal stability results are summarized in Table 1. For these analyses, there were notable differences in community dynamics. Sink A exhibited markedly higher stability than Sink B across all metrics examined. The beta diversity CV was 5-fold lower in Sink A compared to Sink B (CV: 4.9% vs. 26.5%), and the mean squared successive difference was 33-fold lower (MSSD: 0.002 vs. 0.068), indicating substantially reduced volatility in Sink A’s community trajectory (Fig. 2a). Autocorrelation analysis demonstrated significant temporal structure in Sink A (lag-1 autocorrelation: 0.386; Ljung-Box test: χ² = 19.68, p = 0.001), signifying that community states at one timepoint were predictive of subsequent states. In contrast, Sink B showed no significant temporal autocorrelation (lag-1 autocorrelation: −0.207; Ljung-Box test: χ² = 4.17, p = 0.526), suggesting that community fluctuations were relatively stochastic (Fig. 2b).

**Figure 2.**
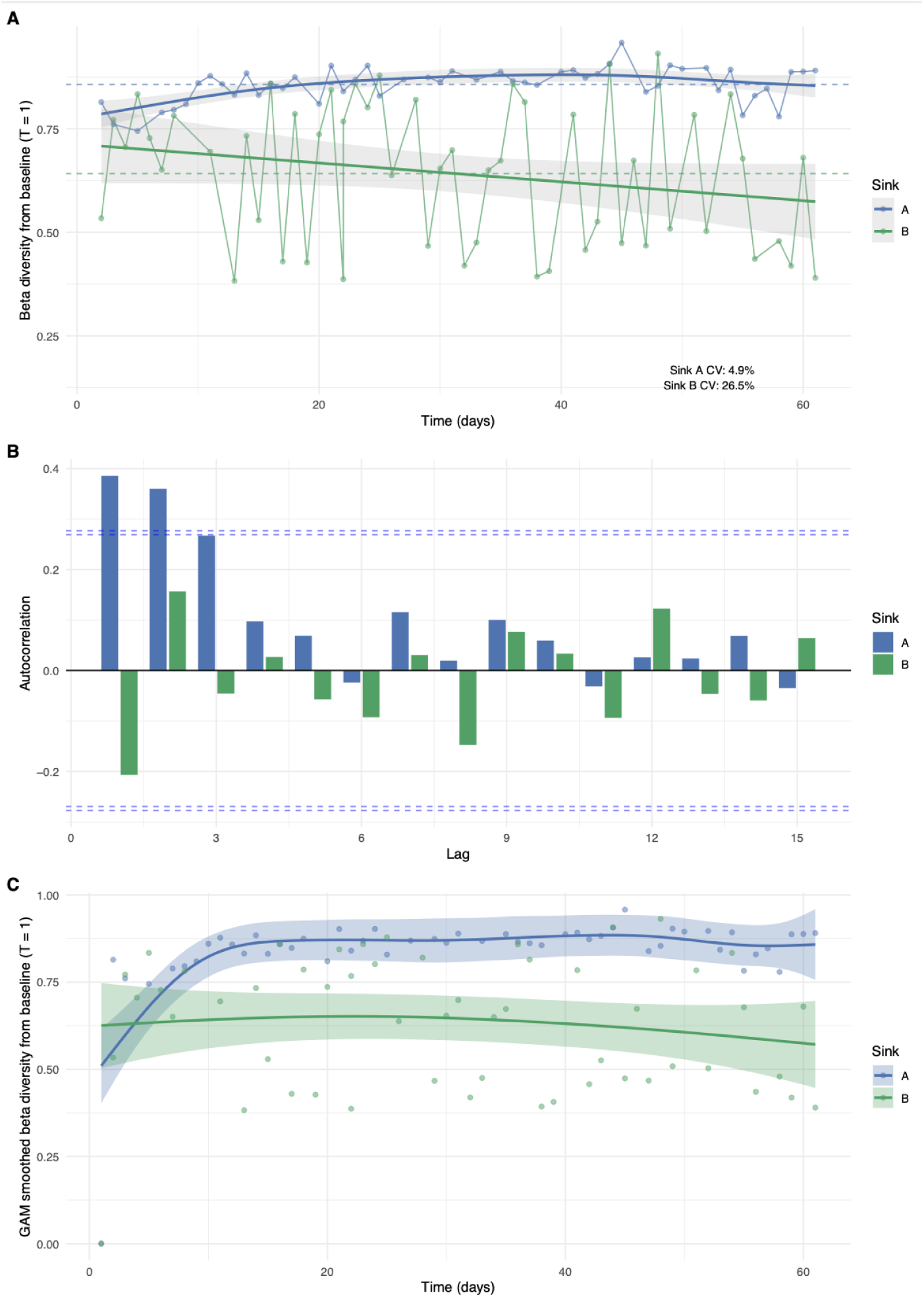
Temporal stability analysis of sink microbial communities. (A) Beta diversity (Bray-Curtis dissimilarity) from the baseline community (T = 1) over time over time for Sink A (blue) and Sink B (green). Solid lines connect consecutive timepoints; dashed horizontal lines indicate mean beta diversity for each sink. Shaded regions indicate 95% confidence intervals. (B) Autocorrelation function (ACF) analysis showing temporal dependence for each sink. Dashed blue lines indicate 95% confidence intervals; bars exceeding these thresholds represent significant autocorrelation. (C) GAM-smoothed temporal trends with 95% confidence intervals.

**Table 1.** Temporal stability metrics for sink microbial communities. Summary statistics comparing the temporal dynamics of Sink A and Sink B, including measures of beta diversity variability (coefficient of variation (CV), mean squared successive difference), temporal autocorrelation (Ljung-Box test), and predictability assessed via generalized additive models (GAM) and autoregressive integrated moving average (ARIMA) modeling with leave-future-out cross-validation.

| Metric | Sink A | Sink B | Interpretation |
| --- | --- | --- | --- |
| N Timepoints | 50 | 53 | Sample size |
| Mean Beta Diversity | 0.857 | 0.6418 | Higher values indicate greater divergence from initial community |
| SD Beta Diversity | 0.0417 | 0.1701 | Measure of spread around the mean |
| CV (%) | 4.8699 | 26.4985 | Sink A (4.87%) is 5x more stable than Sink B (26.50%) |
| Min Beta Diversity | 0.7448 | 0.3827 | Lower bound of community change |
| Max Beta Diversity | 0.9578 | 0.9316 | Upper bound of community change |
| Range | 0.2129 | 0.5489 | Sink A (0.21) has much narrower range than Sink B (0.55) |
| MSSD | 0.0021 | 0.0684 | Sink A (0.002) is 33x less volatile than Sink B (0.068) |
| Lag-1 Autocorrelation | 0.3857 | -0.2068 | Positive values indicate predictability from previous states |
| Ljung-Box Chi-squared | 19.6845 | 4.1651 | Higher values suggest stronger temporal patterns |
| Ljung-Box p-value | 0.0014 | 0.5259 | Sink A shows significant autocorrelation; Sink B does not |
| GAM Deviance Explained (%) | 49.9143 | 3.5363 | Time explains 50% of variance in Sink A but only 3.5% in Sink B |
| GAM Effective df | 5.0863 | 1.5552 | Complexity of the fitted smooth curve |
| **GAM Smooth p-value** | 0.0001 | 0.5411 | Sink A has significant temporal trend; Sink B does not |
| **Best ARIMA Model** | ARIMA(0,1,1) | ARIMA(1,0,0) | Model structure capturing temporal dynamics |
| **ARIMA AIC** | -179.3218 | -34.784 | Lower AIC indicates better model fit |
| **ARIMA RMSE** | 0.0367 | 0.1646 | Lower RMSE indicates better prediction accuracy |
| **ARIMA MAE** | 0.0291 | 0.1448 | Lower MAE indicates better prediction accuracy |
| **CV-RMSE** | 0.0535 | 0.1842 | Number of timepoints used for validation |
| **CV Predictions (n)** | 15 | 16 | Sink A (0.053) has 3.4x lower error than Sink B (0.184) |

Generalized additive models (GAMs) further supported these findings. For Sink A, time explained 49.9% of the deviance in beta diversity, with the smooth term highly significant (p = 5.6 × 10⁻⁵). For Sink B, time explained only 3.5% of deviance, with no significant temporal trend observed (p = 0.54; Fig. 2c). ARIMA modeling identified an integrated moving average model for Sink A (ARIMA(0,1,1); RMSE = 0.037) and a simple autoregressive model for Sink B (ARIMA(1,0,0); RMSE = 0.165), with Sink A showing 4.5-fold lower in-sample prediction error. Leave-future-out cross-validation confirmed that Sink A’s community dynamics were more predictable than those found in Sink B (CV-RMSE: 0.053 vs. 0.184 for Sink B), with prediction error 3.4-fold lower than Sink B. Collectively, these results show that Sink A maintains a stable, deterministic community trajectory with some temporal prediction, where Sink B has stochastic dynamics characteristic of a more perturbed or variable environment. These patterns are consistent with differential usage patterns between the two sinks and align with the enrichment of mature biofilm-associated anaerobes in Sink B, which might reflect episodic disturbance and regrowth cycles.

For minimum sampling frequency needed to predict bacterial communities, Sink A could be sampled as infrequently as every 3-5 days, based on stability metrics (CV, MSSD) and GAM temporal trend modeling, but the detection of significant temporal autocorrelation would require daily or near-daily sampling. For Sink B, community dynamics were stochastic at all sampling frequencies, with no temporal model achieving predictive accuracy, regardless of sampling density.

Differential abundance analysis revealed distinct bacterial community signatures between the two sink environments (Fig. 3). Sink B was characterized by a significant enrichment of strict anaerobes, including the sulfate-reducing bacterium *Desulfovibrio magneticus* (effect size: 2.1; (32)), the fermentative anaerobe *Pelosinus fermentans* (1.75; (33)), and gut-associated obligate anaerobes *Dysgonomonas alginatilytica* (1.69; (34)) and *Bacteroides thetaiotaomicron* (1.34; (35)). This anaerobic signature suggests the presence of mature, oxygen-depleted biofilm communities within Sink B. Sink B was further distinguished by an enrichment of known biofilm-forming, disinfectant-tolerant taxa, including *Raoultella ornithinolytica* (1.94; (36)), *Aeromonas salmonicida* (1.3; (37, 38)), and *Pseudomonas nitroreducens* (1.26; (39)), as well as several species with documented antimicrobial resistance, such as the intrinsically multidrug-resistant opportunistic pathogen *Chryseobacterium indologenes* (1.14; (40))

**Figure 3.**
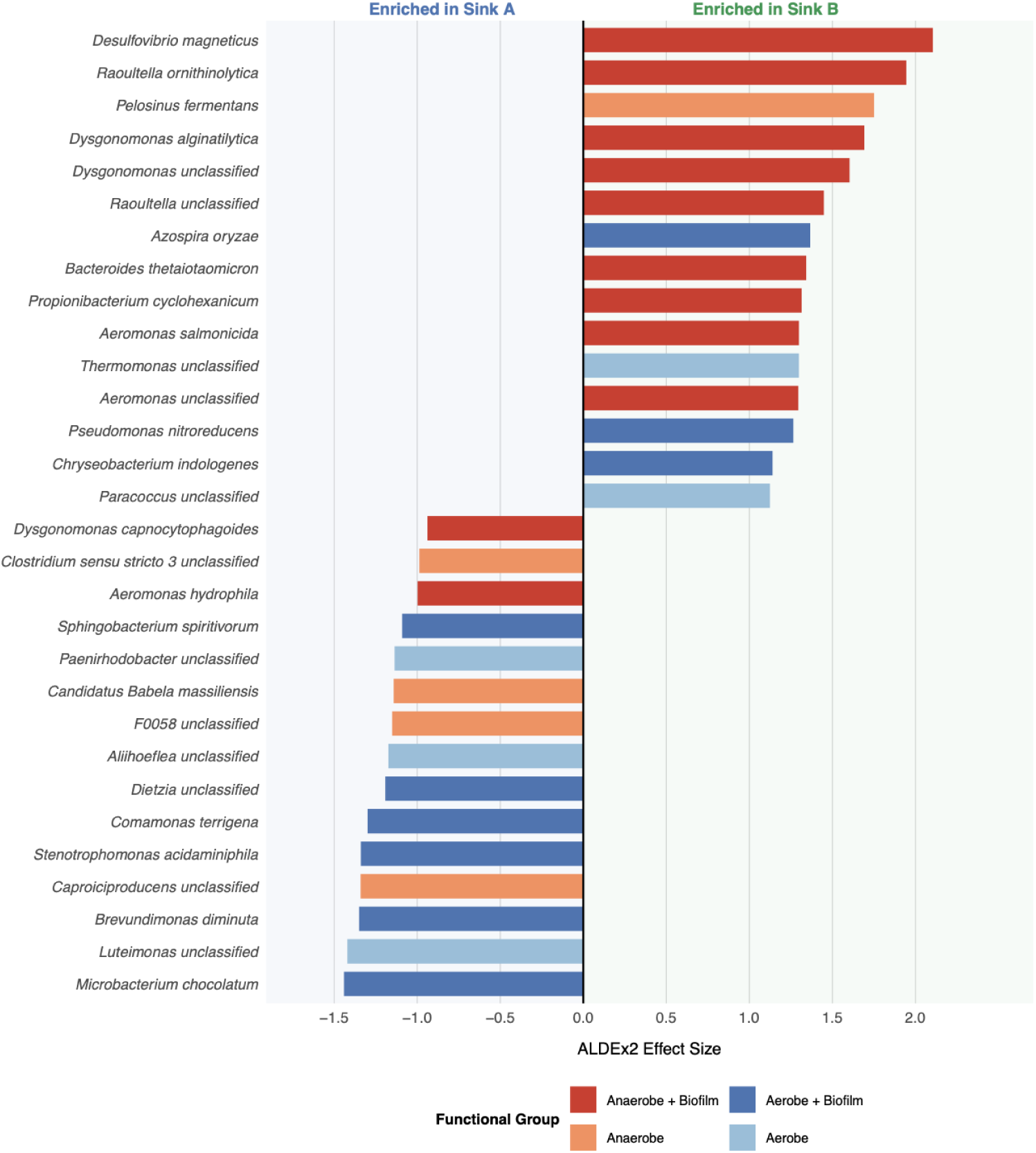
Differential abundance of bacterial taxa between sink environments. ALDEx2 effect sizes for the top 15 differentially abundant species in each sink (n = 30 total). Bars extending left indicate taxa enriched in Sink A; bars extending right indicate taxa enriched in Sink B. Bar colors represent functional classifications: taxa classified as both anaerobic and biofilm-forming (dark red), anaerobic only (orange), both aerobic and biofilm-forming (blue), aerobic only (light blue).

Further, Sink B had an enrichment of oral-associated taxa, suggesting either a greater input from oral hygiene activities or greater survivability of those species upon inoculation. These included oral commensals such as *Streptococcus salivarius* (0.62), *Streptococcus mutans* (0.80), and *Streptococcus parasanguinis* (0.67; (41)); oral anaerobes including *Veillonella dispar* (0.69; (42)) and *Prevotella nigrescens* (0.40; (43)); and upper respiratory commensals such as *Neisseria mucosa* (0.91; (44)) and *Haemophilus parainfluenzae* (0.44; (45)). Sink B was also enriched in preservative-resistant and lipid-degrading taxa, including multiple *Pseudomonas* species that are capable of degrading parabens and other cosmetic preservatives (46), *Methylobacterium-Methylorubrum aquaticum* (0.64; (47, 48)), and lipase-producing bacteria such as *Aeromonas* spp. (49, 50) and *Acinetobacter parvus* (0.46; (51)).

In contrast, Sink A harbored a more aerobic community, with enrichment of strictly aerobic taxa including *Comamonas terrigena* (effect size: -1.3; (52)), *Brevundimonas diminuta* (-1.35; (53)), and *Stenotrophomonas acidaminiphila* (-1.34; (54)), along with skin-associated actinobacteria such as *Dietzia* spp. (-1.19; (55)). Notably, both sinks harbored taxa capable of degrading cosmetic ingredients, though with distinct functional profiles: Sink B was enriched in *P. nitroreducens*, a well-documented degrader of surfactants and parabens (39), while Sink A was enriched in potential cosmetic-degrading taxa including *C. terrigena* (surfactant and aromatic compound degradation; (56, 57)) and *S. acidaminiphila* (xenobiotic degradation; (58)). Specifically, *C. terrigena* is a known degrader of Dialkyl Sulfosuccinates (DASS), which are commonly found in toothpastes and other cosmetics (59). However, *S. acidaminiphila* degrades 4-chloronitrobenzene (4CNB), which is a toxic industrial chemical that is not an ingredient used in personal care items (60). These compositional differences suggest that Sink B maintains conditions favorable for anaerobic biofilm formation and might be subject to greater organic matter accumulation, where Sink A appears to support a more transient, aerobic community with greater contributions from human skin microbiota and environmental water sources.

## Discussion

Although the two sinks were located within the same bathroom and shared identical plumbing characteristics, environmental conditions, and cleaning regimes, we observed variation in alpha diversity over time and distinct bacterial communities (Fig. 1c), suggesting that chemical and physical disturbances associated with individual-specific behaviors altered community structure. Sink A (male) was primarily used for hand washing, toothbrushing, and daily shaving, and Sink B (female) was used for hand washing, toothbrushing, face washing (with cosmetic inputs), and nightly mouthwashing. Therefore, although no antimicrobials were used in the shared bathroom products (hand soap, toothpaste, and cleaners), Sink B had more frequent exposure to biocidal agents, including face wash and mouthwash, a pattern that was reflected in our temporal stability models.

Sink A exhibited characteristics of an equilibrium-state community: low variability (CV = 4.9%), significant temporal autocorrelation, and predictable trajectories that could be modeled with nearly 50% deviance explained by time alone. In contrast, Sink B had high volatility (CV = 26.5%), no significant autocorrelation, and community states that were relatively unpredictable (Fig. 2). The 33-fold difference in mean squared successive difference between sinks further underscores the stochastic dynamics of Sink B, relative to Sink A. Notably, these stability patterns align with our differential abundance findings (Fig. 3). Sink B harbored a greater abundance of strict anaerobes and mature biofilm-associated taxa (*Desulfovibrio*, *Dysgonomonas*, *Bacteroides*), which could reflect cycles of biofilm accumulation, potentially driven by episodic cleaning or variable usage intensity. Such disturbance-recovery dynamics would promote the enrichment of biofilm-forming microbes that are capable of rapid regrowth following perturbation (61). The predictability of Sink A suggests that stable sink environments are more amenable to bacterial monitoring and intervention strategies, with sampling every 3-5 days being sufficient to observe broad stability patterns and daily sampling to resolve fine-scale temporal autocorrelation structure. However, the stochastic nature of Sink B communities might necessitate more frequent sampling to capture representative community states (Fig. 2).

Differential abundance analysis revealed that the two sink environments harbor compositionally and functionally distinct microbial communities, reflecting divergent ecological conditions and usage patterns. The pronounced enrichment of strict anaerobes in Sink B, including sulfate-reducing bacteria (*D. magneticus*), fermentative anaerobes (*P. fermentans*), and obligate anaerobic gut commensals (*Dysgonomonas* spp., *B. thetaiotaomicron*) (32–35), indicates the presence of mature, oxygen-depleted biofilm niches, within the P-trap where organic matter accumulates and oxygen diffusion is limited. This anaerobic signature was accompanied by an enrichment of robust biofilm-forming taxa (*R. ornithinolytica*, *Aeromonas* spp., *P. nitroreducens*) (36–39), consistent with a well-established biofilm consortium capable of withstanding periodic disturbance (Fig. 3; (6)). These results might reflect repeated exposure to mouthwash (e.g., ethanol, quaternary ammonium compounds, and essential oils), which can produce daily antimicrobial exposure that suppresses surface aerobes.

The oral microbiome signal in Sink B, including the enrichment of *Streptococcus*, *Veillonella*, *Prevotella*, and *Neisseria* species (41–45), suggests that this sink either has substantial input from oral hygiene, such as toothbrushing, flossing, and mouthwash use, or that the features of the resident community promotes the survivability of those bacteria upon inoculation (5). Furthermore, the enrichment of preservative-resistant and lipid-degrading taxa (*Pseudomonas*, *Aeromonas*, *Acinetobacter*, *Chryseobacterium*) in Sink B implies potential functional adaptation to personal care products and ingredients found in cosmetics, including parabens, surfactants, and lipids commonly introduced during face washing (Fig. 3; (46–51)).

In contrast, Sink A maintained a more aerobic community dominated by water- and surface-associated taxa (*Brevundimonas*, *Comamonas*, *Stenotrophomonas*) and skin commensals (*Dietzia* spp.) (52–55), suggesting a transient community shaped by hand washing activities and frequent water flow that limits biofilm maturation (Fig. 3). Notably, Sink A was also enriched in *C. terrigena* which degrades surfactant and aromatic compounds that are commonly found in toothpastes and other cosmetics (56, 57), suggesting that sink environments can select for microbial taxa adapted to occupant-specific chemical inputs.

Overall, these compositional differences have potential implications for pathogen persistence and antimicrobial resistance dissemination, as the biofilm-rich, anaerobic environment of Sink B could provide refugia for opportunistic pathogens and facilitate horizontal gene transfer among resistant organisms (10, 28, 62). Understanding how usage patterns shape sink microbiomes can inform targeted cleaning interventions and infrastructure design to minimize reservoirs of clinically relevant bacteria in residential and healthcare settings. Our findings suggest that disturbance regimes likely reduce our ability to predict temporal changes in hospital P-trap microbiomes, consistent with the stochastic dynamics observed in Sink B. Repeated antimicrobial exposure and episodic nutrient pulses might promote cycles of biofilm disruption and rapid regrowth, favoring disturbance-tolerant, biofilm-forming taxa while diminishing temporal autocorrelation. Under these conditions, community composition may fluctuate between semi-stable states, limiting the explanatory power of time-based or deterministic models. However, predictability could increase in hospital sink environments that experience consistent use patterns, such as staff-only hand washing sinks with standardized cleaning schedules and limited organic waste input. In these settings, strong and repeated selective pressures could constrain community composition, leading to convergence on resilient biofilm consortia dominated by antimicrobial-tolerant and anaerobic taxa. Such systems might exhibit increased functional stability, making them more amenable to predictive modeling despite high disturbance frequency.

Future directions could include switching sink usage halfway through the study period to assess whether bacterial community structure shifts toward the baseline associated with each occupant’s usage patterns, converges on an alternative stable state, or if the persistence of biofilms limits community restructuring. Additionally, metagenomic analyses would provide higher-resolution, species- and strain-level taxonomic characterization, enable inclusion of fungal taxa, and allow for the quantification of functional traits, including those associated with biofilm formation, antimicrobial resistance, and virulence. Incorporating genes linked to pathogenicity would facilitate the development of a more comprehensive microbial risk assessment.

## Data Availability

Raw sequence files and metadata are publicly available in the National Center for Biotechnology Information (NCBI) Sequence Read Archive (SRA), under BioProject ID: PRJNA1363208. Additionally, preprocessing and analysis scripts are available on GitHub at https://github.com/hillms/residential_p-traps.

## Acknowledgements

This work was supported primarily by the Engineering Research Centers Program of the National Science Foundation under NSF Cooperative Agreement No. EEC-2133504.

## Conflict of Interest

The authors have no conflicts of interest to declare.

